# Assessing the physiological S-nitrosoproteome reveals nitric oxide–mediated regulatory networks in rice

**DOI:** 10.64898/2026.07.30.741409

**Authors:** Surupa Chakraborty, Sanket Roy, Susmita Poddar, Ankita Choudhuri, Surajit Bhattacharya, Rajib Sengupta

## Abstract

Nitric oxide metabolism-based protein post-translational modifications, such as reversible S-nitrosylation, have been at the ‘pinnacle’ of plant redox research owing to their significant correlation with seed dormancy, interaction with other signaling molecules, plant development and metabolism, biotic and abiotic plant stress responses, immune defense responses against plant pathogens, and senescence. The rapid interconversion of reactive nitrogen species, the abrogation of nitric oxide homeostasis by exogenous supplementation of NO donors and scavengers, the lack of spatio-temporal specificity of NO signaling, and the limited bioavailability or assay sensitivity for detection often limit the effectiveness of identifying and characterizing S-nitrosothiols in plants. Hitherto unknown, we report the first experimental evidence of the total *in vivo* S-nitrosoproteome in *Oryza sativa* L. subsp. *indica*, comprising 134 PSNOs, enriched with 169 putative sites susceptible to S-nitrosylation, without any exogenous supplementation of NO donors. In the present study, mercuric salt-driven facile decomposition of S-nitrosoproteins in the presence of nitrone spin trap 5,5- dimethyl-1-pyrroline N-oxide, resulting in the synthesis of DMPO-nitrone adducts with PSNO-derived protein thiyl radicals in *O. sativa*, has been demonstrated as an efficient and novel strategy for characterizing the PSNOs using mass spectrometry analysis. The evidence of physiological levels of PSNOs was further re-examined in a bi-directional qualitative and quantitative approach involving the 2,3-diaminonaphthalene assay in tandem with fluorescence-based visualization and fluorometric quantification. *In silico* analyses, involving both functional enrichment and pathway prediction analyses, have furthermore revealed unique protein-protein interaction networks and signaling pathways among the S-nitrosoproteome candidates and their predictable physiological roles in *O. sativa indica*, awaiting further *in vitro* validation for their functional correlation in response to S-nitrosylation. In conclusion, the present study provides novel evidence of nitric oxide signaling in rice cultivars under physiological conditions, bringing new insights into the potential *in vivo* transnitrosylation of regulatory or active-site cysteine thiols.

## 1. Introduction

The consequences of exposure to nitric oxide (NO) and its higher redox forms (free radicals, including NO^•^, NO_2_ ^•^, and the nonradical derivatives, including NO^−^, NO^+^, NO Cl, N O , N_2_O_4_, ONOO^−^, ROONO, NO_2_ ^−^, NO_3_ ^−^, and low-molecular-weight S-nitrosothiols (RSNOs), such as *S*-nitrosocysteine (CysSNO), *S*-nitrosohomocysteine (HCysSNO), or *S*- nitrosoglutathione (GSNO)) in plants, range from plant stress response, disease response, seed germination, pollen and pollen tube growth, plant growth and development, fruit ripening, nodule formation, heavy metal stress tolerance, disease resistance, drought and flood stress, oxidative stress responses, gene regulation, effects on phytohormones, and cell signaling effects *via* protein post-translational modifications, such as reversible S-nitrosylation and irreversible tyrosine nitration as well as covalent modifications of fatty acids and nucleotides [1–9]. The source of NO in photosynthetic organisms has been debated by several (in)direct pieces of evidence and assumptions, including lack of L-arginine-dependent NO production *via* nitric oxide synthase (NOS) enzymes in plants, except for *Ostreococcus tauri* and *O. lucimarinus*, unicellular green algae, and *Synechococcus* PCC 7335, a non-heterocystous N_2_-fixing cyanobacterium harboring OtNOS and SyNOS, respectively, NO derived from exogenous supplementation of polyamines (spermine and spermidine) and/or NO donor-mediated NO release, nitrite-dependent NO generation *via* nitrate reductase (NR), utilizing NADH as a cofactor and nitrite (NO_2_ ^−^) and nitrate (NO_3_ ^−^) as competing low- and high-affinity substrates, respectively, evidence of a distinct root-localized plasma membrane-bound nitrite/NO reductase (Ni-NOR) in direct NO production, and possibility of xanthine oxidoreductase (XOR/XDH/XO) pathway as an additional source of endogenous NO *via* nitrite reduction [5, 10–14].

While the mechanistic findings on the source of NO production, uptake, and assimilation in plants have remained unexplained and inconclusive, the biological significance of NO and its redox forms has been largely underscored by the evidence of NO-mediated S-nitrosylation in *Arabidopsis thaliana*, *Brassica juncea*, *Nicotiana tabacum*, *N. tabacum Bright Yellow-2 cells, N. plumbaginifolia*, *Pisum sativum*, *Solanum tuberosum*, *S. habrochaites, S. lycopersicum*, *Kalanchoe pinnata*, *Helianthus annuus, Populus trichocarpa*, *Oryza sativa,* and the unicellular green alga, *Micrasterias denticulata* [1,6,15–27]. Despite the evidence of RSNOs, mainly S-nitrosoproteins (PSNOs), in several plant varieties, including *O. sativa*, studies on rice plant-derived PSNOs have been limited to forward genetic screening of *Os noe1*, *Os noe1-2* mutants, overexpression (GSNOR-O *noe1*) or knockdown (GSNOR-R) transgenic plants, with perturbed (S)NO homeostasis, and *OSNIA1*, *OSNIA2*, and *OSNOA1*, and *CPL1* gene correlation with NO production in *rhs1* knockout mutants, whereby estimation of *in vivo* PSNO content was hindered by their exposure to a physiological excess of NO donor(s) such as sodium nitroprusside (SNP) and CysSNO, NO scavenger, and/or NOS inhibitor [25–27]. Although these findings represented the implications of an abrogated NO homeostasis in a wider landscape of rice plants, the evidence of basal or physiological levels of thiol-based nitrosative modification, i.e., S-nitrosylation, has been an unresolved enigma due to i) a lack of identification and characterization of the total *O. sativa* proteome, ii) experimental pitfalls from the widely used, Biotin-Switch Assay (BSA) technique for PSNO/ RSNO quantification, ii) lower sensitivity of detection from Saville assay-like techniques, iii) use of exogenous addition of NO or exposure of whole cell lysates to physiological or synthetic NO donors that might overestimate the number of -SNO functions on protein thiols, iv) long-term exposure to NO gas or any other exogenous NO source that might rewire the endogenous NO production, v) prolonged proteomic analysis of this ‘reversible’ modification, and vi) possibilities of PSNO formation or degradation during the sample preparation upon influence of additional factors such as, changes in pH, heat, light, proximity to endogenous NOS-like enzymes and S-denitrosylating systems, chances of oxidation of cellular thiols, right choice of growth media lacking any prior nitrogenous or NO source, presence of transition metal ions or complexes, and presence of other biological artifacts.

Given the key challenges in PSNO detection, specifically methodological limitations, and considering the scope of *in vivo* S-nitrosylation in *O. sativa indica*, we aimed to investigate the physiological concentrations of PSNOs without any NO donor treatment. The latter being relatively stable (BDE of the S-N bond usually ranges between 23.3 and 32.4 kcal/mol [28]), identification and characterization of the candidates in the S-nitrosoproteome in rice (*indica* cultivar) using an improved proteomic technique involving a bi-directional qualitative and quantitative approach has been another major focus of this study.

## 2. Experimental Methodology

### 2.1 Rice seed germination, seedling growth and crude cell-free root extract preparation

An adequate quantity of *Oryza sativa* L. (IR64) seeds was selected and subjected to surface sterilization following to the protocol of Das et. al (2017) [29]. The surface-sterilized seeds were aseptically placed on the solidified water agar medium and incubated under suitable growth conditions for germination. 14th-day-old seedlings were collected, and their root tissues were carefully excised for subsequent extraction for experimental analyses. After that, 600 mg of dry weight of root samples were frozen with liquid nitrogen and powdered using a mortar and pestle. Thereafter, the powdered root extract was dissolved in Chelex 100-treated 0.1 M pre-chilled phosphate buffer (pH 7.4; 3 × 1 mL), containing 1 mM EDTA, and reconstituted to homogeneity. Following cold centrifugation (10000 rpm for 5 min at 4 °C) of the freshly prepared root extracts (200 mg/ml), the pellets containing cell debris were sedimented, and the supernatants were carefully transferred, aliquoted into several vials, and stored at -20 °C for further use. The lysate protein content in three biological replicates ranged from 0.125 to 1.2 mg/ml (batch-to-batch variability), as determined spectrophotometrically using the Bradford reagent at 595 nm on a spectrophotometer.

### 2.2 Quantification of PSNOs with L-CysSNO as a standard

The cell-free protein extracts from the freshly excised *O. sativa* root samples were initially treated with 60% perchloric acid (PCA; 9.2 M) to a final concentration of 0.5 M at 4 °C for 5 min, and the homogeneous mixture was centrifuged (7000 rpm for 10 min at 4 °C) and fractionated into a deproteinized supernatant and protein-enriched insoluble pellets. The supernatants containing low-molecular-weight components and any interfering metabolites were discarded, and the pellets were reconstituted in 0.1 M phosphate buffer (pH 7.4), containing 1 mM EDTA, followed by their resuspension in the 2,3-diaminonaphthalene (DAN; (Tokyo Chemical Industry Pvt. Ltd. (TCI)) assay reaction mixture (0.1 M HCl, 0.5 mM DAN, 0.5 mM HgCl_2_). S-nitroso functions on the rice plant proteins were quantified following an Hg^2+^-catalyzed decomposition of PSNOs to protein thiolate (PS^−^) and NO^+^, with the concomitant detection of naphthotriazole (NAT), a fluorescent triazole derivative of DAN, when the released NO^+^ reacts with DAN at an acidic pH. The aliquots of the lysates were treated with a final concentration of 100 μM each of DAN and HgCl_2_, and the reaction solution was incubated in the dark for 10 min at room temperature. To prevent prolonged incubation and chances of any false positives, the DAN assay reaction in the final solution in each reaction set was thereafter terminated with 0.4 M NaOH [30]. PSNO calibration was performed with a comparable standard curve derived from freshly prepared standard solutions of S-nitroso-L-cysteine (L-CysSNO). In positive control experiments, the aliquots were treated with reduced glutathione (5 mM final working solution) and assessed for residual PSNOs following the same DAN assay-based quantification. All measurements of PSNOs were carried out using a spectrofluorometer (Hitachi F-4600 Fluorescence Spectrophotometer) (λ_ex_ = 375 nm, λ_em_ = 450 nm; excitation and emission slits = 5 nm; photomultiplier voltage = 700 V).

### 2.3 Preparation of standard RSNO solutions

L-CysSNO was used as a standard S-nitrosothiol for PSNO quantification. L-CysSNO solution was freshly prepared by mixing the acidified (0.1 M HCl) stock solutions of 100 mM Cysteine and NaNO_2,_ and further diluting it (1:1) with pre-chilled phosphate buffer as described previously. The prepared final stock was incubated on ice for 30 min, and its concentration was assessed by a Hitachi U2910 UV-vis spectrophotometer. The molar extinction coefficient at 334 nm is ε_CysSNO_ = 594 M^-1^ cm^-1^ [31].

### 2.4 Quantification of protein thiols (PSH)

Free thiol (-SH) concentration of proteins from *O. sativa* root samples was determined by the 5,5′-dithio-bis (2-nitrobenzoic acid) or DTNB (Ellman’s reagent; Sisco Research Laboratories Pvt. Ltd. (SRL)) assay using a Hitachi U2910 UV-vis spectrophotometer at room temperature. Crude cell-free protein extracts were obtained following a 60% PCA (9.2 M)-based deproteinization method. The insoluble fractionates or pellets were reconstituted in 0.5-2 mM DTNB final working solution (100 mM DTNB dissolved in ethanol (solubility 8 mg/ml), 0.1 M potassium phosphate pH 7.4, 1 mM EDTA; 10 ml) on ice and the final reaction volume (700 µl) was analyzed by the colorimetric detection of the 2-nitro-5- thiobenzoate anion (TNB^2-^) at 412 nm, formed as a by-product during the reaction, which corresponded to the -SH function per mg of protein. The spectrophotometric detection was determined using an optical quartz absorption cuvette (1000 µl chamber volume) with a wavelength scan performed within the visible spectral range (400-700 nm) at a scan speed of 1200 nm/min. An absorbance maximum at 412 nm was attributed to the concentration of free thiol function in protein cell lysates, whereby the molar extinction coefficient of TNB^2-^ at 412 nm is 13600 M^-1^ cm^-1^ (maximum possible absorbance = 0.9 at 412 nm corresponds to 66.17 µM RSH) [32, 33].

### 2.5 Fluorescence microscopy

Thin transverse sections of root tissues were promptly transferred with a brush and rinsed with pre-chilled phosphate buffer in a Petri dish until analysis. Thereafter, 3-4 plant sections per reaction set were transferred into 1.5 ml microfuge tubes, treated with 0.1 M phosphate buffer, and then incubated with 100 µM DAN and HgCl_2_ reaction mix (1:1 molar concentration) at room temperature (25 °C) for 10 minutes in a dark chamber. The assay reactions were terminated by the addition of 1 M NaOH to a total reaction volume of 100 µl. The samples were transferred from the reaction solution onto clean, sterilized slides and then visualized under coverslips using an Olympus fluorescence microscope, equipped with an Olympus DP28 detector (DTT sets; Figure 2a) and a Leica DM6 B Fluorescence microscope; software LAS X (Leica Microsystems for GSH sets: Figure 2b). The bright-field (DIC), DAPI channel images, and autofluorescence observations were performed precisely and uniformly for all samples to prevent any chances of photobleaching from overexposure. In 1,4-dithiothreitol or Cleland’s reagent (DTT)-treated and reduced glutathione (GSH; HIMEDIA Laboratories Pvt. Ltd)-treated sets, samples were additionally incubated for 10 min at 25 °C with 5 mM DTT and GSH, respectively, prior to DAN-HgCl_2_ treatment, as already mentioned. Data were obtained by visualizing the fluorescence intensities with excitation and emission maxima of 358 nm and 461 nm, respectively. Affixation of cells onto slides with chemical fixatives was avoided to prevent any assay interference in visualizing the basal level of PSNOs. Image acquisition was performed with a Leica microscope equipped with a camera. Images were processed with ImageJ (https://imagej.net/ij/) without obscuring any original data.

**Fig. 1:**
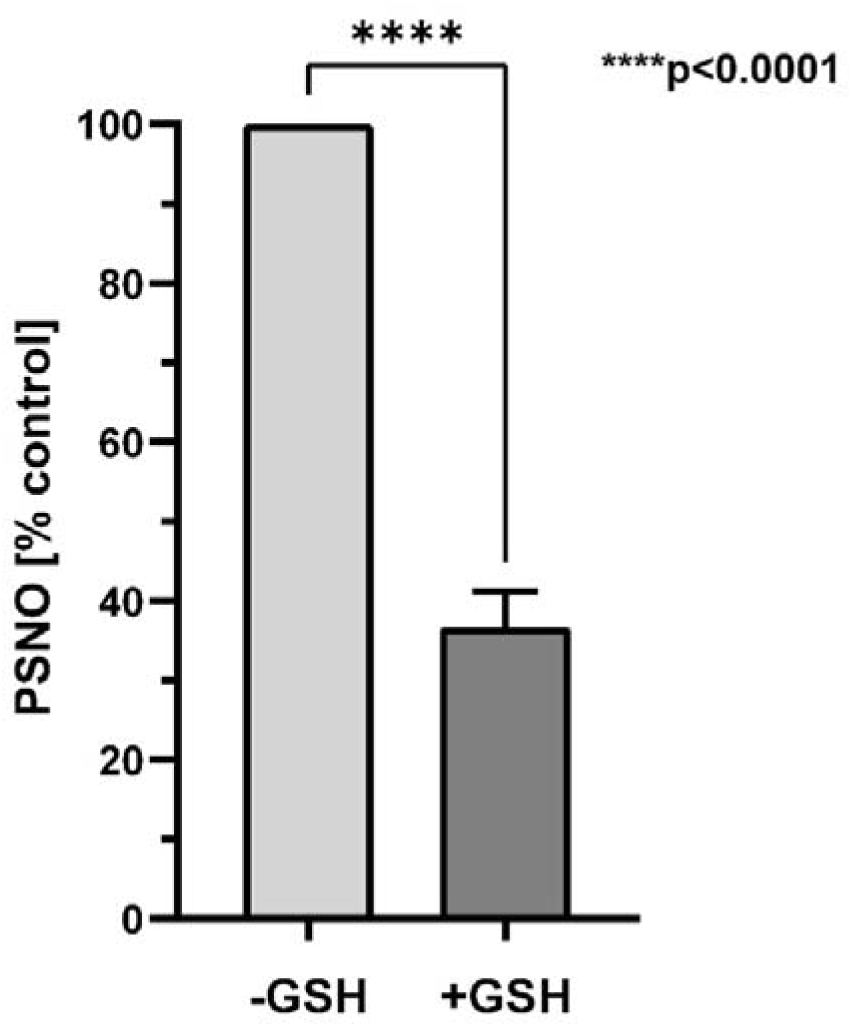
GSH-catalyzed S-denitrosylation of *O. sativa*-derived PSNOs. Experiments were carried out in Chelex 100-treated 0.1 M potassium phosphate buffer (pH 7.4), containing 1 mM EDTA at RT (25 °C). Freshly prepared 100 mM GSH stocks were used at an optimal concentration of 5 mM. PSNO and PSH quantification were performed utilizing the DAN and DTNB assays, respectively, as summarized in the Experimental Methodology. The data are expressed as mean values of two independent biological replicates, each with two technical replicates, with SD (*n* = 4).

**Fig. 2a:**
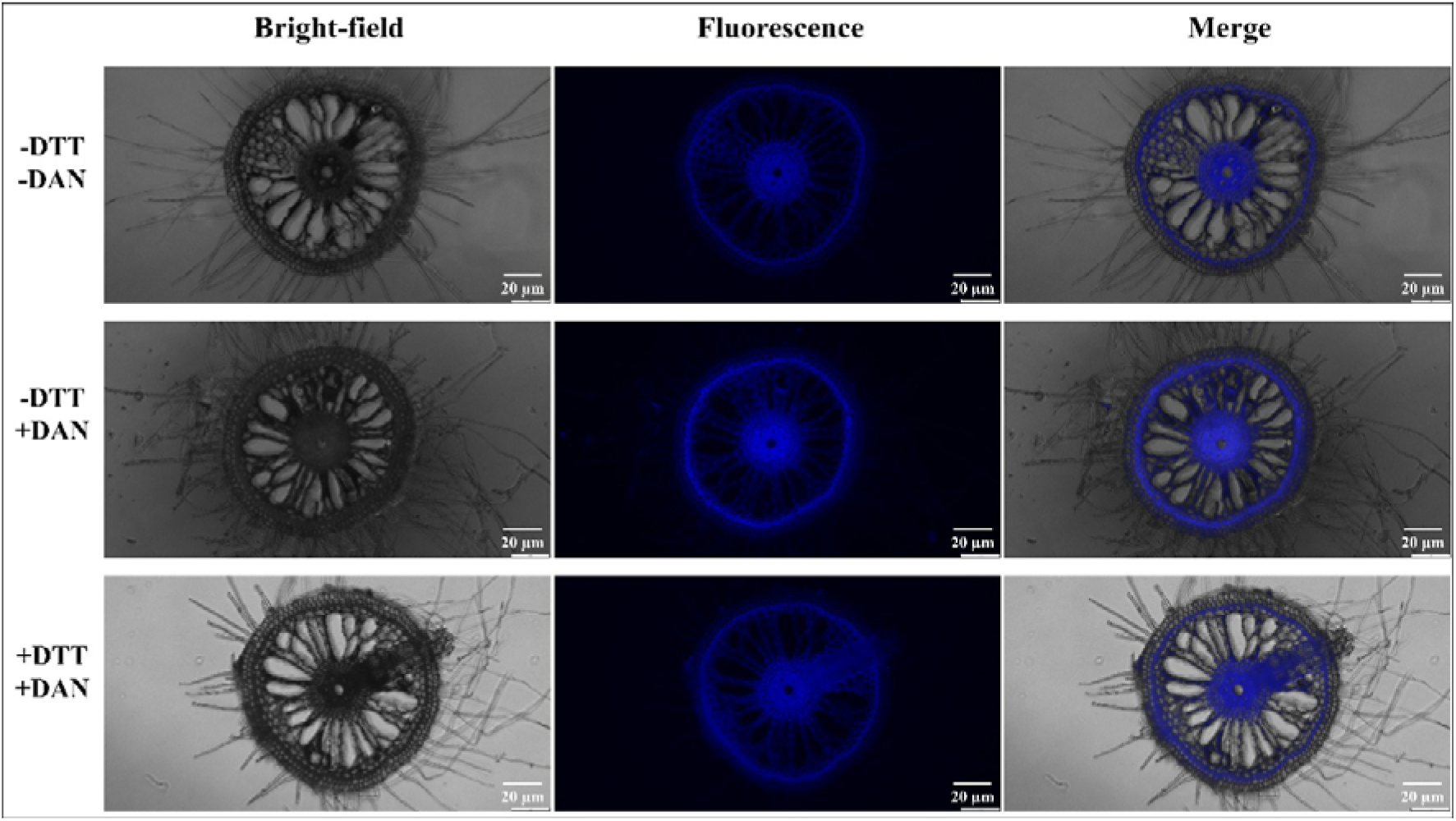
DTT catabolizes the *in vivo* S-nitrosoproteome in *O. sativa* root tip cells. S-nitrosothiols were determined by the DAN assay in tandem with fluorescence microscopy imaging, as elaborated in Experimental Methodology. (a) Untreated root tip cells were used as controls to obtain representative autofluorescence images. DAN- and Hg^2+^-treated intracellular PSNOs were recorded without (panel 2) and with treatment of 5 mM DTT (panel 3). Panel images of root tip cells have been acquired under bright-field and blue fluorescence conditions at 20X magnification, with a 20 µm scale bar. The blue fluorescence intensity of the DAN-treated sets, relative to that of the autofluorescence (untreated) sets, represents the fluorescence emitted upon releasing NAT.

**Fig. 2b:**
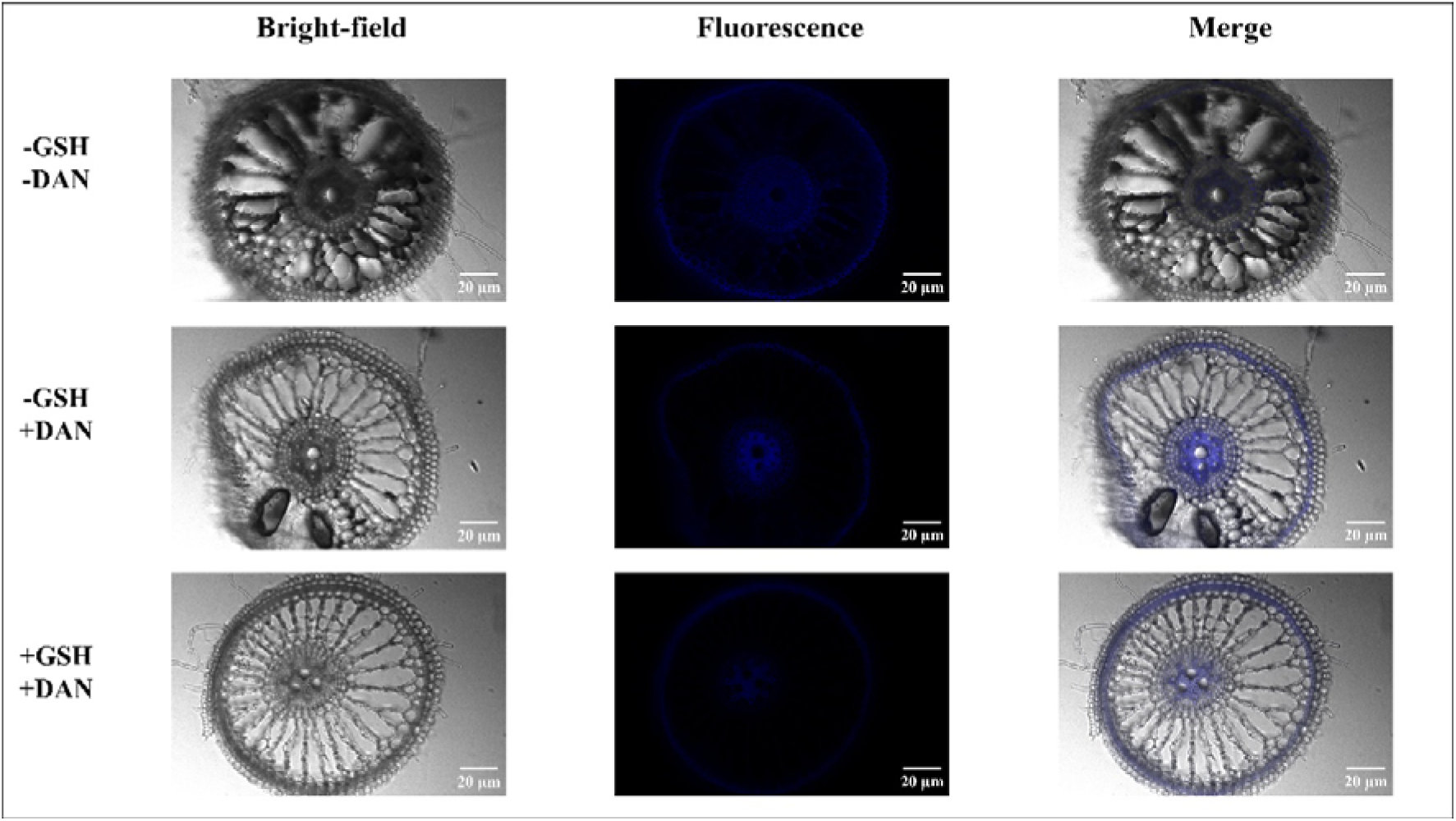
GSH catalyzes S-denitrosylation of the *in vivo* S-nitrosoproteome in *O. sativa* root tip cells. S-nitrosothiols were determined by the DAN assay in tandem with fluorescence microscopy imaging, as elaborated in Experimental Methodology. (a) Untreated root tip cells were used as controls to obtain representative autofluorescence images. DAN- and Hg^2+^- treated intracellular PSNOs were recorded without (panel 2) and with treatment of 5 mM GSH (panel 3). Panel images of root tip cells have been acquired under bright-field and blue fluorescence conditions at 20X magnification, with a 20 µm scale bar. The blue fluorescence intensity of the DAN-treated sets, relative to that of the autofluorescence (untreated) sets, represents the fluorescence emitted upon releasing NAT.

### 2.6 Synthesis of PS-DMPO nitrone and mass spectrometry using LC-ESI-MS/MS

Protein thiyl radicals (PS^•^) were produced by the chemical decomposition (homolysis) of endogenous PSNOs, as previously reported [34]. In brief, a 50 mM stock solution of 5,5- dimethyl-1-pyrroline N-oxide (DMPO) (5 mg/ml; Sigma-Aldrich) in dimethyl sulfoxide (DMSO) and 50 mM HgCl2 were added to 1 ml of 0.1 M HCl to achieve a final concentration of 10 mM. The incubation of the final reaction mix containing DMPO and HgCl_2_ was continued at RT for a few minutes. Subsequently, fresh cell lysates (1.2 mg/ml) were supplemented with 1 mM DMPO-HgCl_2_ solution and incubated for 30 min at 4 °C in a dark chamber. The resulting lysate enriched with PS-DMPO nitrone adducts was treated with 1X Trypsin-EDTA solution (10X Trypsin-EDTA solution (HIMEDIA), followed by an overnight incubation at RT. Thereafter, the 100 ng/ul final sample was lyophilized under ultralow-temperature (−110°C cold trap) and vacuum conditions using a LaboGene ScanVac CoolSafe freeze dryer. The resulting hygroscopic solid content in the microfuge tube was reconstituted in 5 μl of 0.1% formic acid and analyzed by liquid chromatography-high-resolution tandem mass spectrometry with electrospray ionization (Orbitrap). The sample was injected into an Easy nLC 1000 LTQ Orbitrap XL system (Thermo Fisher Scientific) in ESI (+) ion mode, equipped with an Easy Spray C18 nanocolumn, using water and acetonitrile as standards. The mass spectrum was acquired with a resolution of 60000 for full MS over the m/z range of 300-2000, at a sample injection flow rate of 0.3 µl/min, with a total run time of 120 min. The total protein database of *O. sativa indica* was retrieved from UniProt, and the DMPO modification at the respective cysteine residues was provided to the pipeline of the Proteome Discoverer 1.4 software (Thermo Fisher Scientific) for the identification of DMPO- conjugated peptides. The search parameters were restricted to a maximum of 2 missed cleavage sites, while the limits of mass tolerances of the singly charged precursor ion to be processed were set to 10 ppm for MS ions and 0.8 Da for MS/MS fragment ions.

### 2.7 Gene Ontology (GO) analysis and Protein-protein interaction study of the identified PSNOs

In order to identify the biological and molecular functions of the identified PSNOs, gene ontology analysis was performed using ShinyGO v0.77: Gene Ontology Enrichment Analysis (https://bioinformatics.sdstate.edu/go77/) webserver. The STRING database (https://string-db.org/) has been used to study the protein-protein interaction network of the identified proteins in the *O. sativa indica* cultivar.

### 2.8 Statistical methods

Statistical significance of PSNO and PSH content (nmol/mg) derived from rice root extract samples was determined using an unpaired *t*-test, performed in GraphPad Prism 8.0.2 software (GraphPad Software Inc., San Diego, CA). Results are obtained from independent biological and technical replicates (n=4 to 6, as mentioned in the legends) and reported as ±SD.

## 3 Results

### 3.1 Comparative profiling of PSNOs and PSH in O. sativa

Freshly prepared cell-free lysates from rice seedlings cultured in 0.8% water agar, devoid of any NO donor supplementation preceding or concurrent with the lysis procedure, revealed a novel evidence of a basal level of PSNOs in *O. sativa*. The procedure to quantify endogenous PSNO concentrations in rice root extracts, as described in Materials and Methods, revealed S-nitrosoproteins (1.52 ± 0.08 nmol/mg) with high purity [Table 1]. Additionally, any possibility of transition metal ion-catalyzed decomposition of PSNOs, which could potentially abrogate the accurate quantification of PSNOs, was minimized by utilizing Chelex100-enriched chilled phosphate buffer, containing 1 mM EDTA, during the analysis. To discriminate the endogenous populations of S-nitrosylated (−S-NO) and reduced or free (−S-H) thiols corresponding to the total proteome of *O. sativa*, fluorometric (DAN assay) and spectrophotometric (DTNB assay) approaches were concomitantly carried out from the same batch of extracts, revealing consistently comparable results (1.52 ± 0.08 vs 50.29 ± 4.42 nanomoles per milligram of protein, respectively) [Table 1]. Within the scope of this reactive thiol-based protein post-translational modification, we were interested in assessing the stability of the *O. sativa* S-nitrosoproteome in the presence of physiological concentrations of reduced glutathione. Interestingly, incubation of *O. sativa-*derived PSNOs (*Os*PSNO)- enriched deproteinized pellets (retentate) with phosphate buffer containing 5 mM GSH for 10 min at room temperature, followed by the standard DAN assay procedure, resulted in efficient catabolism, i.e., S-denitrosylation of PSNOs, leading to a 60-70% decrease in fluorescence intensity as compared to the control set, without GSH treatment. The residual stable PSNOs accounted for about ∼35-40% (36.7 ± 4.5 in percentage; *n=*4) [Figure 1] as compared to the control sets for each batch (without GSH treatment; normalized to 100%), which confirms the specificity of the reversible S-nitrosylation mechanism and the susceptibility of a majority of *Os*PSNOs to the physiological reductant, GSH.

**Table 1:** Fluorometric and spectrophotometric determination of *in vivo* S-nitrosoproteins (PSNO) and reduced protein thiol (PSH) concentrations, respectively, as a function of endogenous NO and its higher redox species in root extracts of *O. sativa indica.* The data are expressed as mean values of three independent biological replicates, each with two technical replicates, with SD (*n* = 6).

| Sample | Content (nmol/mg) |
| --- | --- |
| PSNO concentration | 1.52 ± 0.08 |
| PSH concentration | 50.29 ± 4.42 |

### 3.2 Visualization of DTT- and GSH-catalyzed S-denitrosylation of PSNOs in O. sativa root transverse sections

While the evidence for significant endogenous levels of PSNOs has been quantitatively demonstrated, suggesting their possible *in vivo* biochemical, structural, and functional significance, we sought to validate the novelty of our experimental findings using a qualitative approach. In parallel studies of fluorescence microscopy, ultrathin rice root transverse sections were observed in the DAN-HgCl_2_-treated sets [Figures 2(d-l)] and analyzed for their fluorescence intensities, corresponding to the fluorogenic NAT released in the vicinity of the intracellular PSNOs as compared to the untreated control (DAN-HgCl_2_- lacking sets) [Figures 2(a-b)], exhibiting autofluorescence of the cell walls. The excitation and emission wavelengths of the released fluorophore, NAT, are 375 nm and 450 nm, respectively. Utilizing the intrinsic, highly fluorescent property of NAT, upon reaction of endogenous PSNOs with HgCl_2_, the relative change in fluorescence intensities was observed in the external layers (epidermis, exodermis, sclerenchyma), aerenchyma, metaxylem, cortex, and stele regions. With DAN-HgCl_2_ exposure, the localization and relative concentration of PSNOs can be determined, owing to their significantly higher fluorescence intensities, as observed by the DAPI and merged channel images [Figures 2(d-f)]. The fluorescence intensities corresponded to the intracellular presence of PSNOs in O. sativa, suggesting that the native sulfhydryl (thiol) moieties of susceptible proteins undergo in vivo S-nitrosylation without prior exposure to NO donors. At sub-optimal concentrations (100 µM) of DAN and HgCl_2_, and after precise incubation with the samples for 10 min in the dark, neither reagent altered the structural integrity of the cells, as evidenced in [Figures 2(a-l)]. A rapid catabolism of intracellular PSNOs was concomitantly observed, resulting in a significantly reduced fluorescence upon exogenous supplementation with the synthetic reductant, DTT (5 mM), and the physiological reductant, GSH (5 mM) [Figures 2(g-i) and 2(j-l)], as described in Materials and Methods.

### 3.3 Identification and characterization of DMPO-conjugated PSNOs identified by LC-ESI-MS/MS

The PS-DMPO nitrone adducts that were formed by mercuric salt (HgCl_2_)-catalyzed decomposition of *in vivo* PSNOs in the presence of DMPO were further analyzed by nano LC-ESI-MS/MS. The nitrone trap conjugated thiyl adducts were detected in mass spectral analysis as their dynamic modification on the cleaved peptide sequences bearing susceptible cysteine residues (+*m/z* 111.068 Da), leading to the identification of 134 PSNOs, in the molecular mass ranges of 4.5 to 567.4 kDa, with 169 DMPO binding sites [Table 2]. The adducts were mostly detected as their [M+H]^+^, i.e., protonated, monoisotopic mass of the peptides, and the MS/MS scans were recorded for the most abundant ions, derived from the adjacent raw time-integrated LC-MS chromatogram [Supplementary Figure 1]. Interestingly, in the same mass spectral analysis, the peak positions and relative abundances extracted by the built-in software also revealed evidence of *in vivo* GSH in untreated plant cell extracts. The evidence of the [M+H]^+^ peak corresponding to the GS-DMPO nitrone adduct (*m/z* 419.32, 100% relative abundance) is in good agreement with the studies of Stoyanovsky et al. (1996) [35]. In comparison, the remaining peaks correspond to the peptide-derived thiyl- DMPO nitrone adducts, characterized by relatively lower relative abundances [Supplementary Figure 1].

**Table 2:**
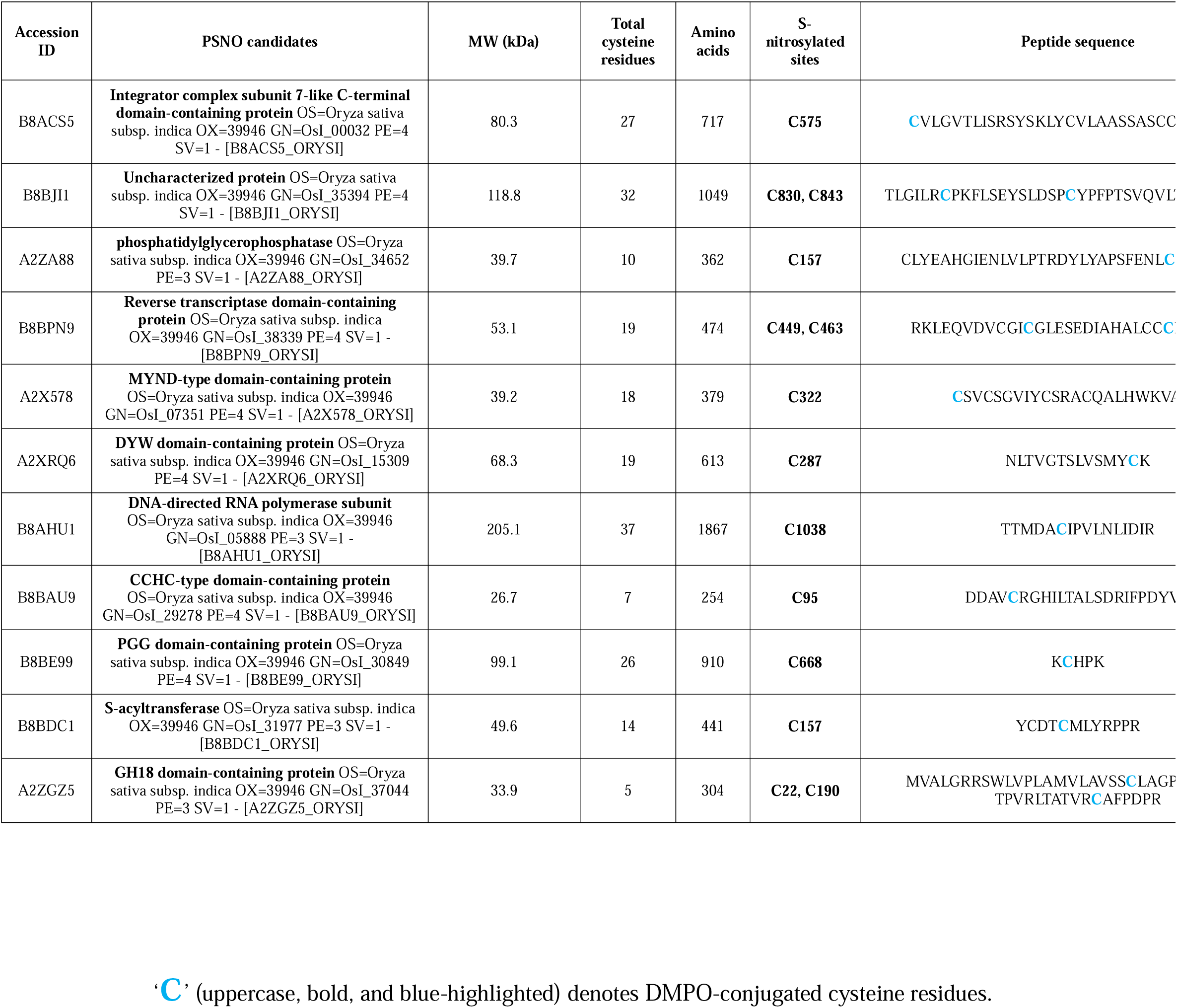
DMPO-conjugated PSNOs identified in *Oryza sativa* L. subsp. *indica* with a score above 0 using LC-ESI-MS/MS.

### 3.4 GO enrichment analysis of identified PSNOs

To gain further insight into the functional significance of the identified endogenous PSNOs, GO enrichment analysis was performed using the ShinyGo web server (ver 0.77). The analysis indicated that the PSNOs are associated with multiple biological processes, predominantly involving RNA metabolism, transcriptional regulation, enzymatic phosphoregulation, and acyltransferase-mediated modification. Among the biological processes, significant enrichment was observed for SnRNA processing, SnRNA metabolism, ncRNA processing, and metabolism involving Integrator complex subunit 7-like C-terminal domain-containing protein (B8ACS5) [Figure 3]. Following that, involvement of Phosphatidylglycerophosphatase (A2ZA88) in protein dephosphorylation and peptidyl-tyrosine dephosphorylation suggested the involvement in reversible signaling pathways.

**Fig. 3:**
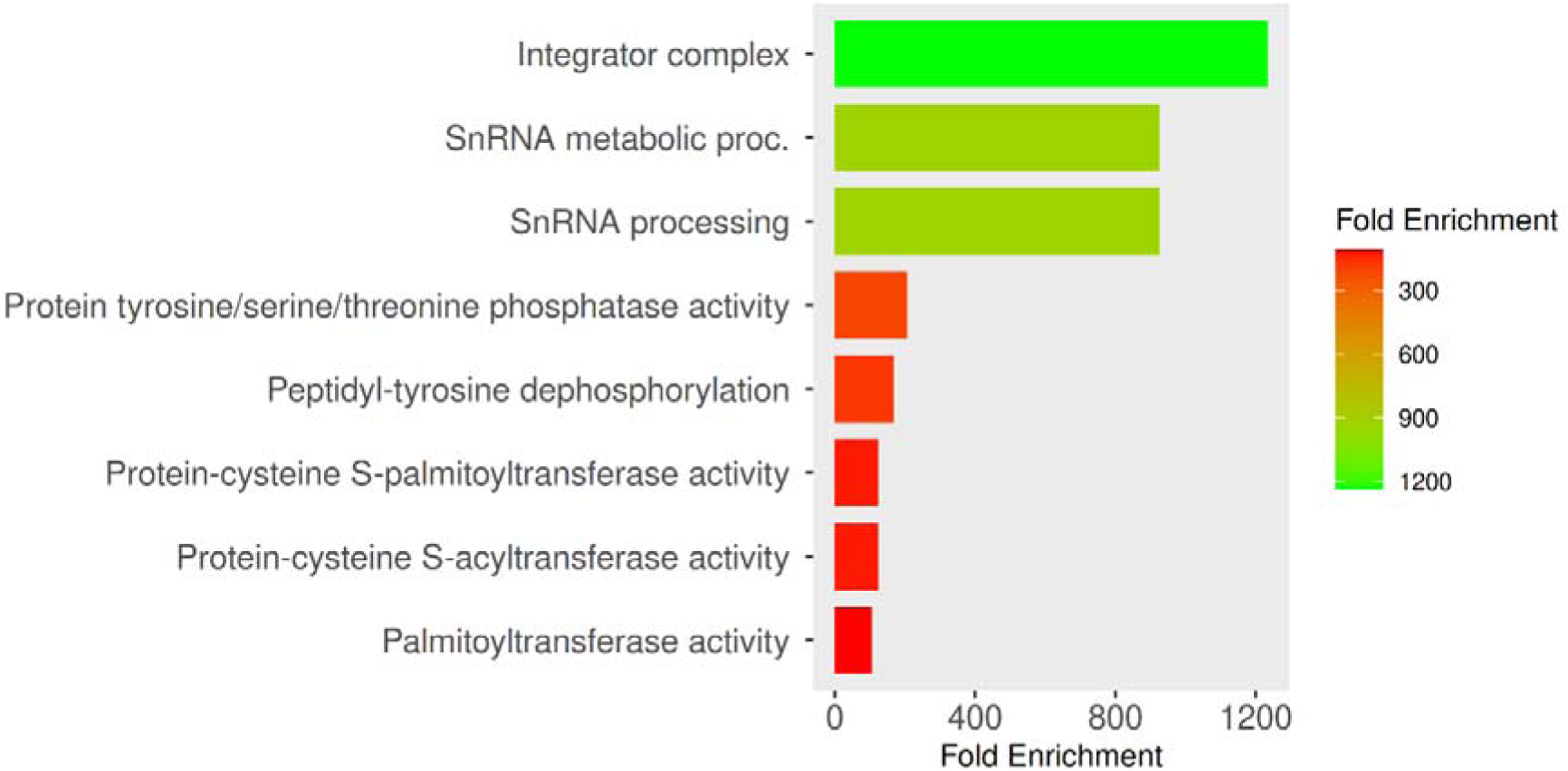
GO enrichment analysis of the identified PSNO dataset with a score above 0. The magnitude of fold enrichment of significantly enriched GO terms is represented by horizontal bars of varying lengths and a (red to green) color gradient, with increasing green intensities corresponding to a higher enrichment.

Apart from that, phosphatase-associated terms such as protein tyrosine/serine/threonine phosphatase activity and phosphoprotein phosphatase activity were significantly represented. Another major molecular function consisted of palmitoyltransferase activity, S-acyltransferase activity, protein-cysteine S-palmitoyltransferase activity, and protein-cysteine S-acyltransferase activity, represented by S-acyltransferase (B8BDC1), implying coordination between redox signaling and lipid-based protein modification. GO terms related to DNA-directed RNA polymerase activity and nucleotidyltransferase activity were enriched through the DNA-directed RNA polymerase subunit protein (B8AHU1), suggesting that components of the transcriptional machinery are potential targets of S-nitrosylation. In addition, zinc ion binding and transition metal ion binding were enriched through DYW domain-containing protein (A2XRQ6) and CCHC-type domain-containing protein (B8BAU9), implying the presence of metal-dependent catalytic or nucleic acid-binding proteins within the identified PSNO repertoire [Figure 3].

### 3.5 Protein-protein interaction network analysis of identified PSNOs

To further unravel the biological interactome among the identified S-nitrosylated proteins, a putative protein-protein interaction network analysis was performed using the STRING database. The resulting network demonstrated sparse but biologically relevant interactions, elucidating that instead of clustered in a highly connected network complex, the identified PSNOs belong to a diverse and functional pathways, suggesting the independent role in various biological pathways. In the obtained protein-protein interaction network, only one notable interaction cluster was observed between the Uncharacterized protein B8ACS5 and the DNA-directed RNA polymerase subunit [Figure 4].

**Fig. 4:**
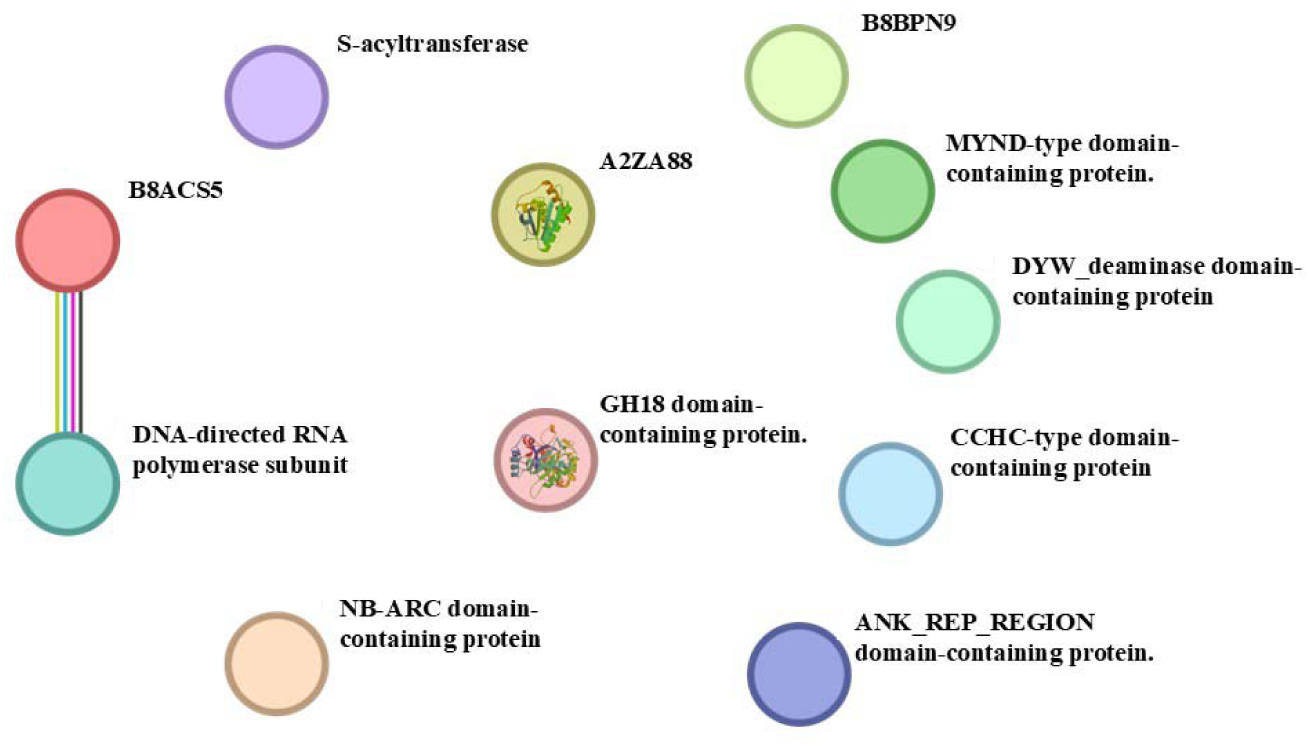
Protein-protein interaction network of *O. sativa indica-*derived PSNOs generated using STRING database among 11 analyzed PSNOs (MS score > 0).

In the preliminary STRING protein-protein interaction analysis with the total *O. sativa* S-nitrosoproteome, encompassing 134 PSNO candidates, the association network consisted of 134 nodes or proteins in the network, 42 edges (42 known and 29 predicted interactions among 134 proteins), with an average node degree of 0.627, an average local clustering coefficient of 0.194, and a PPI enrichment p-value of 0.0136 (< 0.05), implying high statistical significance [Supplementary Figure 2]. Functional enrichment analysis of the entire submitted dataset revealed grouped statistical enrichment observations for a plethora of functions and sub-functions. The results indicate an enrichment of identified genes or their characteristic proteins in the network that are predominantly associated with gene regulatory processes, such as post-transcriptional gene silencing by RNA (5 proteins with a false discovery rate (FDR) of 0.0139) processes and a broader category of gene silencing by RNA (6 proteins with an FDR of 0.0477).

## 4 Discussion

In the present study, experimental evidence of *in vivo* S-nitrosylation of proteins and their reversible decomposition or S-denitrosylation catalyzed by GSH has been presented in the *O. sativa indica* model system. Notably, a major fraction (60-65%) of the *Os*PSNOs with a molecular mass range of 4.5 to 567.4 kDa were substrates of GSH, a physiological non-enzymatic S-denitrosylating agent [36–38]. S-denitrosylation of PSNOs by GSH reflects the differential susceptibility of *in vivo Os*PSNOs towards GSH-dependent decomposition, which can be mechanistically explained by a bimolecular nucleophilic substitution (*S*_N_2) reaction [38]. While copious levels of reduced GSH in leaves and root tip cells have been reported in amounts of 344.40 nmol/g of fresh weight of wild-type *Arabidopsis thaliana*, subcellular compartmentalized GSH has been detected in mitochondria, nuclei, peroxisomes, cytosol, chloroplasts, and vacuoles at 8.7−15.1, 6.4, 2.6−4.8, 2.8−4.5, 1−1.4, and 0.01−0.14 millimolar concentrations, respectively, thus gaining acceptance as the most abundant intracellular thiol in plant systems [39–42]. Thus, the formation of GSNO *via* i) S-nitrosylation of a fraction of GSH [43], ii) nitro-linolenic acid (NO_2_-Ln)-dependent or NR-mediated NO release and its further interaction with GSH [7, 44–47], iii) reaction of GSH with other NO-derived intermediates (S-nitrosylating species), or iv) indirectly *via* formation of low-molecular-weight (LMW) RSNOs and HMW PSNOs, followed by their subsequent rapid trans-S-nitrosylation to GSH [37]. The interaction of protein thiols with acidified nitrite, peroxynitrite, NO, and its higher oxidized species; metal-catalyzed S-nitrosylation; interaction of protein thiyl radicals (PS^•^) with NO, and transnitrosylation of the NO moiety (NO^+^) from GSNO, other cellular S-nitrosylating species (L-CysSNO, HCysSNO), or PSNOs to specific cysteine thiol residues of target proteins might reasonably explain the mass spectrometry-based evidence of 134 *in vivo* PSNO candidates in *O. sativa* [48].

To date, predicted protein-coding sequences for about 37,440 entries, corresponding to 37,430 protein-coding genes, have been deposited in the UniProt Knowledgebase (UniProtKB) (www.uniprot.org/proteomes/UP000007015) for *O. sativa indica* (rice), of which 134 identified PSNOs comprising 46 uncharacterized and 88 characterized candidates could be detected following LC-ESI-MS/MS analysis, which is indicative of the composite S-nitrosoproteome in *Oryza sativa* L. subsp. *indica* [Figure 5]. In contrast to the conventional experimental paradigm that involves exogenous addition of any physiological (CysSNO or GSNO) or non-physiological (sodium nitroprusside (SNP), S-nitroso-N-acetyl-DL-penicillamine (SNAP), nitroglycerin, DEA-NONOate, etc.) NO donors, the qualitative and quantitative data presented herein establish *in vivo* S-nitrosylation of *Os*-derived proteins, surpassing the detectable limits of the assays, without any external NO stimulus. Although empirical, the current study offers a conceptual blueprint for further analysis of the effects of *in vivo* S-nitrosylation on enzymatic activity and the functional consequences of the identified PSNOs in a wider landscape of plant physiology, immunity, and thiol-dependent redox regulation in cells.

**Fig. 5:**
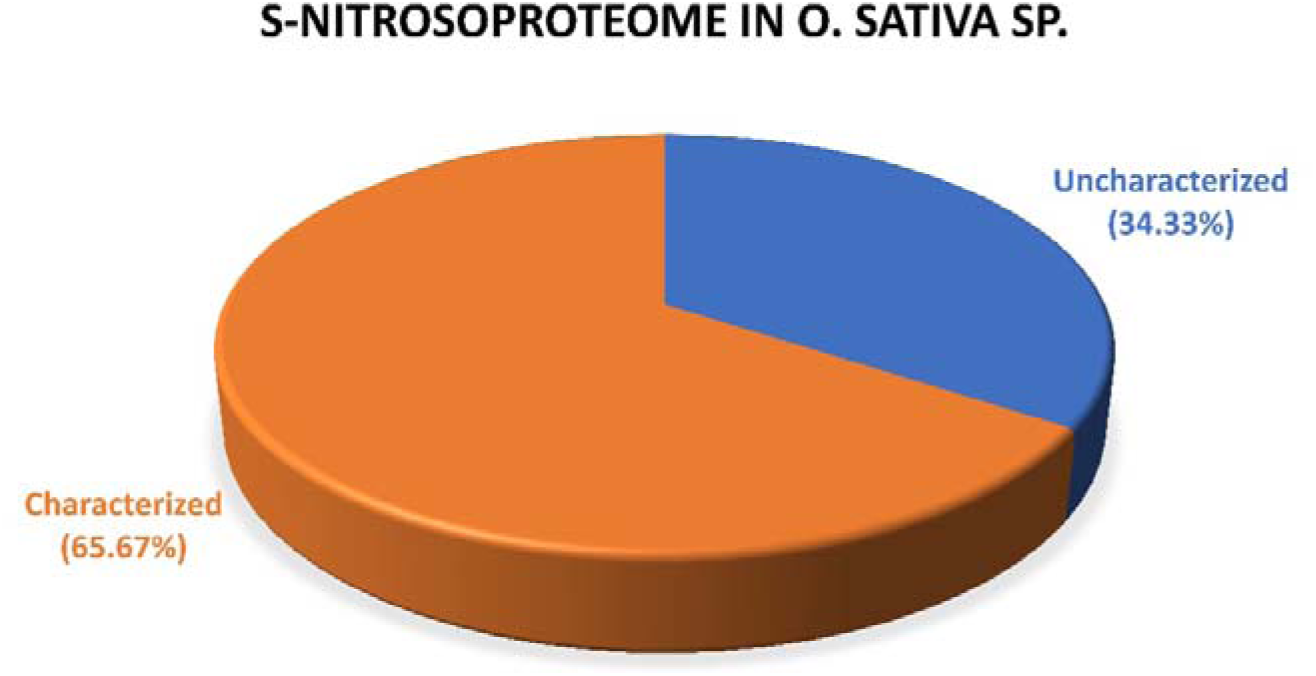
3D pie-chart illustrating the proportion (expressed in percentages) of characterized and uncharacterized protein candidates from the identified *in vivo* S-nitrosoproteome of *O. sativa indica*.

In biological systems, the participation of free (sulfhydryl; −SH) and S-nitrosylated (SNO-modified) thiols in a dynamic redox equilibrium necessitates sensitive assays for their detection in nanomolar amounts per mg of protein. In contrast with mammalian cells, plant cell lysate preparation is significantly more challenging primarily due to the abundant concentrations of interfering secondary metabolites, and pigments, and the presence of a rigid cellulose, hemicellulose, and pectin-rich cell wall that obscures protein extraction at higher yields using a simple lysis buffer. Such methodological hindrances could be overcome by grinding freshly excised rice root samples in liquid nitrogen to a fine powder, following their homogenization with pre-chilled buffer, as mentioned previously. While the DAN and DTNB assays merit quantification of the *in vivo* PSNO and PSH concentrations at nanomolar to picomolar ranges, the presence of a plethora of endogenous LMW compounds, such as phenolic acids, tannins, flavonoids, phytoalexins, and tocopherols, in plant cell-free extracts or lysates makes their quantification overwhelmingly difficult. Because of their abilities to act as quenchers, fluorophores with overlapping excitation and emission spectra, or their direct interference with DAN, the LMW compounds and the perchloric acid-rich resolving supernatant could be separated from protein-enriched fractions, which were precipitated as pellets by a PCA-based deproteinization method, prior to further biochemical analysis [49].

The intramolecular cyclization and aromatization of DAN, leading to the formation of its fluorescent triazole derivative, NAT, follow an HgCl_2_ (Hg^2+^)-mediated selective cleavage of the S−N bond in PSNOs, releasing NO^+^ or its equivalent species, which then interacts with DAN. However, direct evidence of the DAN assay for the formation of NAT in intact cells has only been provided so far in pathogenic and non-pathogenic strains of *Vibrio cholerae,* utilizing the fluorescence property of the released NAT and confocal microscopy [34]. The visualization of S-nitrosylation of protein thiols by microscopy is particularly challenging because of the relatively labile nature of the S–NO bond. To overcome this limitation, specialized techniques have been employed over the past decade that can potentially transform unstable −SNO groups into stable and detectable labels. Thus, with the advent of several robust biochemical PSNO detection techniques, S-nitrosylation-based imaging using fluorescence or confocal microscopy has become a crucial aspect in redox biology. This is commonly accomplished through selective chemical labeling strategies that enable the detection and localization of PSNOs within biological systems. In our previous study [34], we developed a method based on the DAN assay principle to monitor NAT release from PSNOs and subsequently employed confocal microscopy to determine its subcellular localization. In the current study, the DAN assay has proven to be instrumental in the qualitative detection of PSNOs in transverse histological sections of rice roots. Owing to its relatively small molecular size and lipophilicity due to its conjugated naphthalene rings, DAN is easily permeable through the cell wall and the phospholipid bilayer of the plasma membrane of the root tip cells. While the biochemical DAN assay provides the quantitative estimation of *in vivo* PSNO concentration and their stability in the presence of physiological concentrations of GSH, the fluorescence microscopy images show a similar trend, whereby DAN-HgCl_2_-treated samples exhibit relatively higher fluorescence intensities than the autofluorescence sets, lacking any treatment. The images in Panel B of Figures 2a and 2b typically show highly concentrated, bright fluorescence of NAT, especially in the central column (stele) regions of the root transverse sections, including the vascular tissues: xylem and phloem. A moderately high fluorescence intensity is observed in the sclerenchyma (the third layer of concentric cylinders from the external periphery to the internal) of the same panel as compared to that of the autofluorescence sets. Additionally, faint signals were detectable in the root hairs. Consistent with the trend of quantitative biochemical estimation of PSNO content in untreated (lacking DTT or GSH) and GSH-treated samples, Panels B and C of Figures 2a and 2b reveal significant differences in relative fluorescence intensities, implying the S-denitrosylation of PSNOs in intact cells. The inherent background fluorescence images (Panel A of Figures 2a and 2b) corresponded to the autofluorescence values measured biochemically, which were subtracted from each of the DAN-HgCl_2_-treated sets to avoid any consideration of false positives.

To further strengthen our evidence of *in vivo* S-nitrosylation of proteins in *Oryza sativa* L. subsp. *indica*, we extended the scope of PSNO detection from fluorometric quantification to site-specific identification of (−SNO)-modified protein thiols, regardless of their low or high cellular abundances, utilizing an immune-spin trapping agent or redox probe, DMPO. The experimental workflow presented in Figure 6 illustrates the mercuric salt (Hg^2+^)-induced chemolysis of PSNOs, leading to the formation of protein thiol-conjugated DMPO-nitrone adducts, which, after trypsin-mediated cleavage, are utilized for mass spectrometric analysis. Initially established as an EPR-based spin trapping technique with high feasibility of detection of O- and S-centered reactive radical species (R−S^•^, G−S^•^, [Hb-Fe^IV^=O]^•^, [Mb- Fe^IV^=O]^•^, etc.) in model biochemical and proteomic platforms [35, 50, and references thereof], the implications of DMPO for downstream proteomic analysis have now been optimized for the specificity of detection of peptide or protein thiyl radicals (R−S^•^ or P−S^•^; R represents the rest of the molecule, and P represents protein) *via* photolytic or chemolytic homolysis of PSNOs [34, 51, and the current study]. To our knowledge, this is the first direct experimental evidence for the mapping of PSNO candidates of the *in vivo* S-nitrosoproteome in *O. sativa*, harnessing the benefit of the formation of remarkably stable PS−DMPO nitrone adducts [Figure 6], which are subsequently formed from low-stable cyclic nitroxide intermediates (the half-life of GSH-derived GS−DMPO nitroxide is ∼2 min) [50]. The lower final concentrations of DMPO (1 mM) and high specificity of HgCl_2_ (Hg^2+^) in S−N bond decomposition allowed for the *Os*PSNO detection in rice root extracts by the concomitant labeling of only the PSNO-derived protein thiyl radicals with DMPO.

**Fig. 6:**
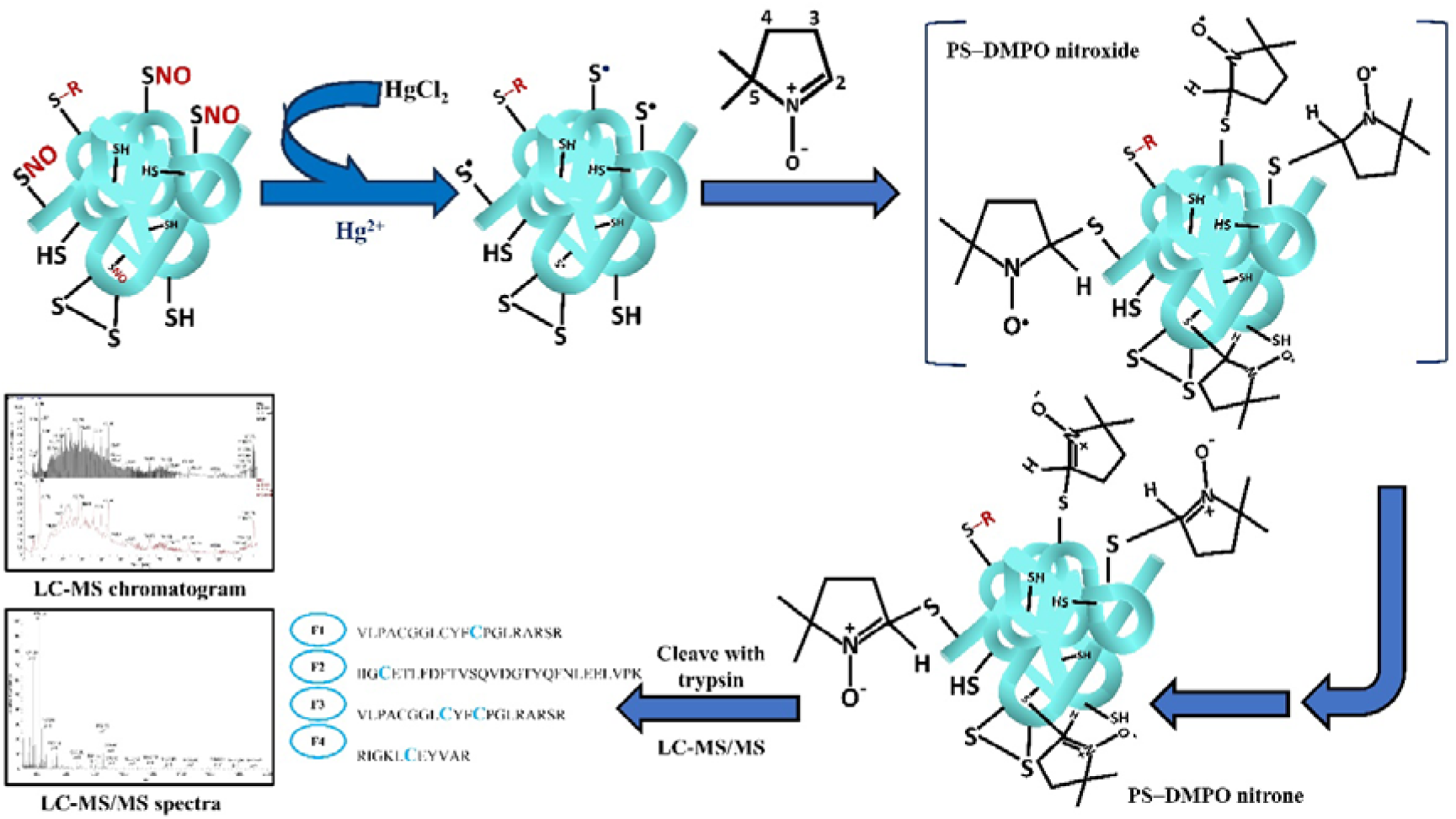
Schematic representation of chemolysis/DMPO-mediated spin trapping of an arbitrary PSNO and identification of its S-nitrosylation sites using tandem mass spectrometry analysis. Mercury salt (Hg^2+^) mediates decomposition or chemolytic cleavage of PSNO(s) derived from purified protein or a complex mixture of proteins, yielding protein thiyl radicals (P−S^•^). The addition of DMPO to this reaction system results in PS-DMPO nitrone adducts, which can be detected as DMPO-bound susceptible cysteine(s) enriched fragments in LC-MS/MS analysis, following trypsin digestion.

The integration of the mass spectrometry (LC-MS/MS)-based proteomic workflow data with GO and STRING enrichment analyses is the gold standard to identify several significantly enriched components, including biological processes, molecular functions, and cellular components, associated with the identified protein dataset (here, identified PSNO candidates). While GO analysis with the proteomic data is inherently challenging with a vast dataset of 134 total identified PSNO candidates, yielding no significant enrichment, the same functional enrichment tool with a significant threshold or FDR cutoff of ≤ 0.05, considering 11 PSNOs (MS score > 0), identified certain major enrichments. Among the chosen dataset, the simultaneous and significant enrichment of the integrator complex, snRNA metabolic process, and snRNA processing as a part of cellular component analysis is consistent with the predominance of RNA metabolism and post-transcriptional regulation observed throughout the functional analyses. Molecular function and biological process analyses localized a significant proportion of identified PSNOs to harboring protein tyrosine/serine/threonine phosphatase activity, peptidyl tyrosine dephosphorylation, protein-cysteine S-palmitoyltransferase activity, protein-cysteine S-acyltransferase activity, and palmitoyltransferase activity. The higher enrichment of the phosphatase and/or palmitoyltransferase functions might be suggestive of the NO- or, specifically, S-nitrosylation-mediated broader modulation of the enzymes and their downstream effects on reversible phosphorylation-dependent signaling and lipid-associated protein dynamics in the cellular milieu. However, the ultimate question of whether there are any functional crosstalks or coordinated regulation of protein activity through S-nitrosylation, phosphorylation, and palmitoylation-like processes underlying the identified proteomic dataset remains elusive and awaits further experimental validation. The STRING analysis further complements the GO analysis data with the observation of physically or functionally interconnected clusters in the entire dataset of 134 PSNOs, with an important advantage of multiple biological pathways associated with the annotated S-nitrosylated proteins, rather than isolated statistical enrichments.

## Credit authorship contribution statement

Experiments were carried out by S.C., S.R., S.P., and A.C. Manuscript writing, methodology designing, data curation, and corrections were carried out by S.C., S.R., S.B., and R.S. R.S. and S.B. conceptualized the original idea and supervised the research work. All authors reviewed and approved the final manuscript.

## Data availability statement

Data is contained within the article.

## Declaration of competing interest

The authors declare that they have no competing financial interests, personal relationships with other people or organizations that could have inappropriately influenced the work, the position presented in, or the review of, the manuscript entitled.

## Acknowledgement

SB acknowledges the research grants CRG/2022/007083 and (2296 (Sanc.)/STBT- 13015/8/2024-WBSCST SEC) from Anusandhan National Research Foundation, Govt. of India, and Department of Science and Technology and Biotechnology, Govt. of West Bengal, respectively. The authors sincerely thank Dr. Amit Ranjan Maity’s lab, Amity University Kolkata and Dr. Avishek Banik’s lab, Institute of Health Sciences, Presidency University (2^nd^ campus), Kolkata for the technical support with the Olympus and Leica DM 6 B fluorescence microscopy imaging facilities, respectively.

---

**Supplementary Fig. 1:**
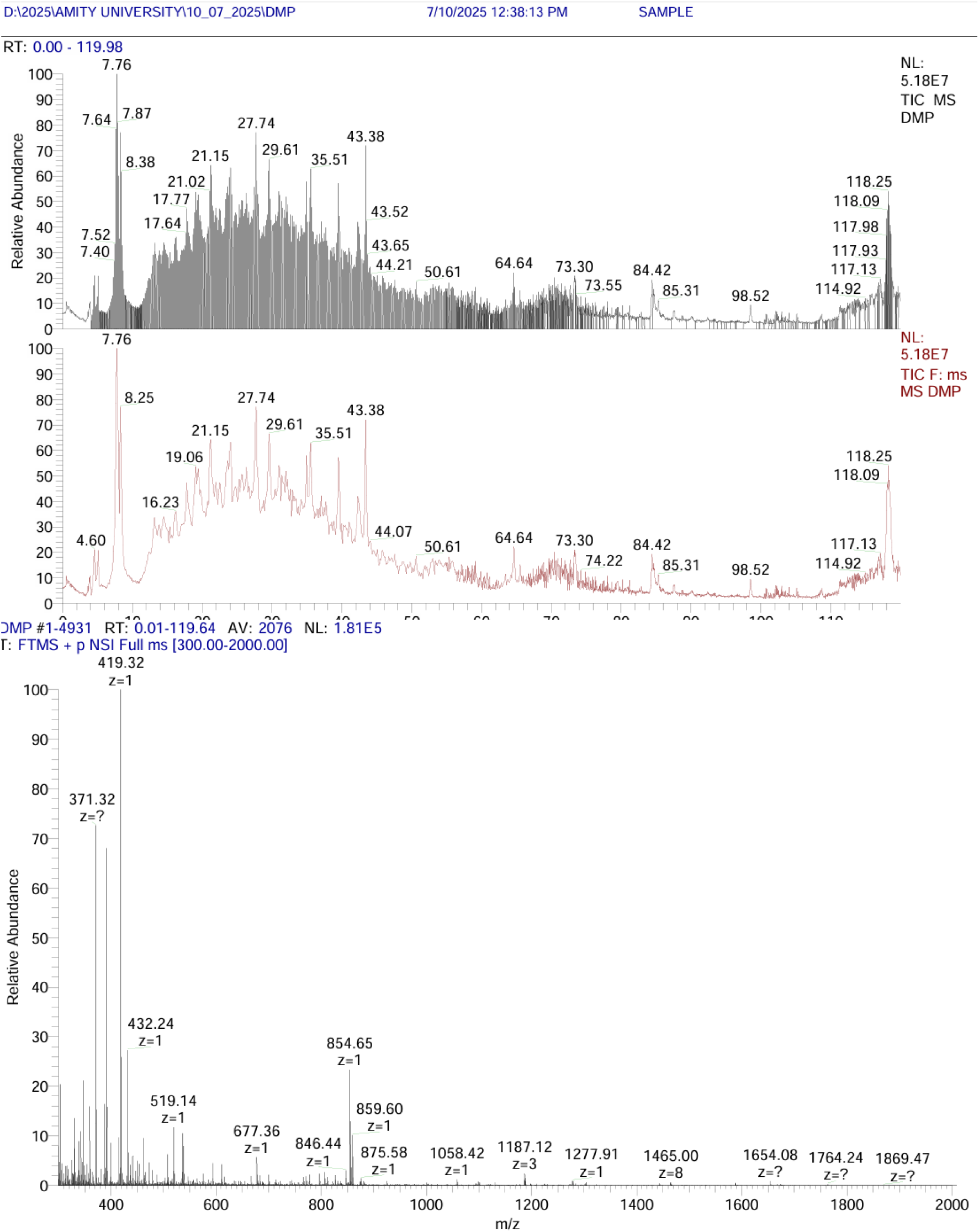
LC-MS and LC-MS/MS chromatograms representing the spectra of the peptides eluted under their respective peaks at given time points. Full MS scan was acquired in an Orbitrap analyzer (Fourier Transform Mass Spectrometer; FTMS), in positive ion mode (p), using NanoSpray Ionization (NSI) over the specified m/z range of 300.00- 2000.00. The maximum relative abundance was attributed to the GS-DMPO nitrone accumulated in *O. sativa* root extracts [35]. Assay conditions are specified in Experimental Methodology.

**Supplementary Fig. 2:**
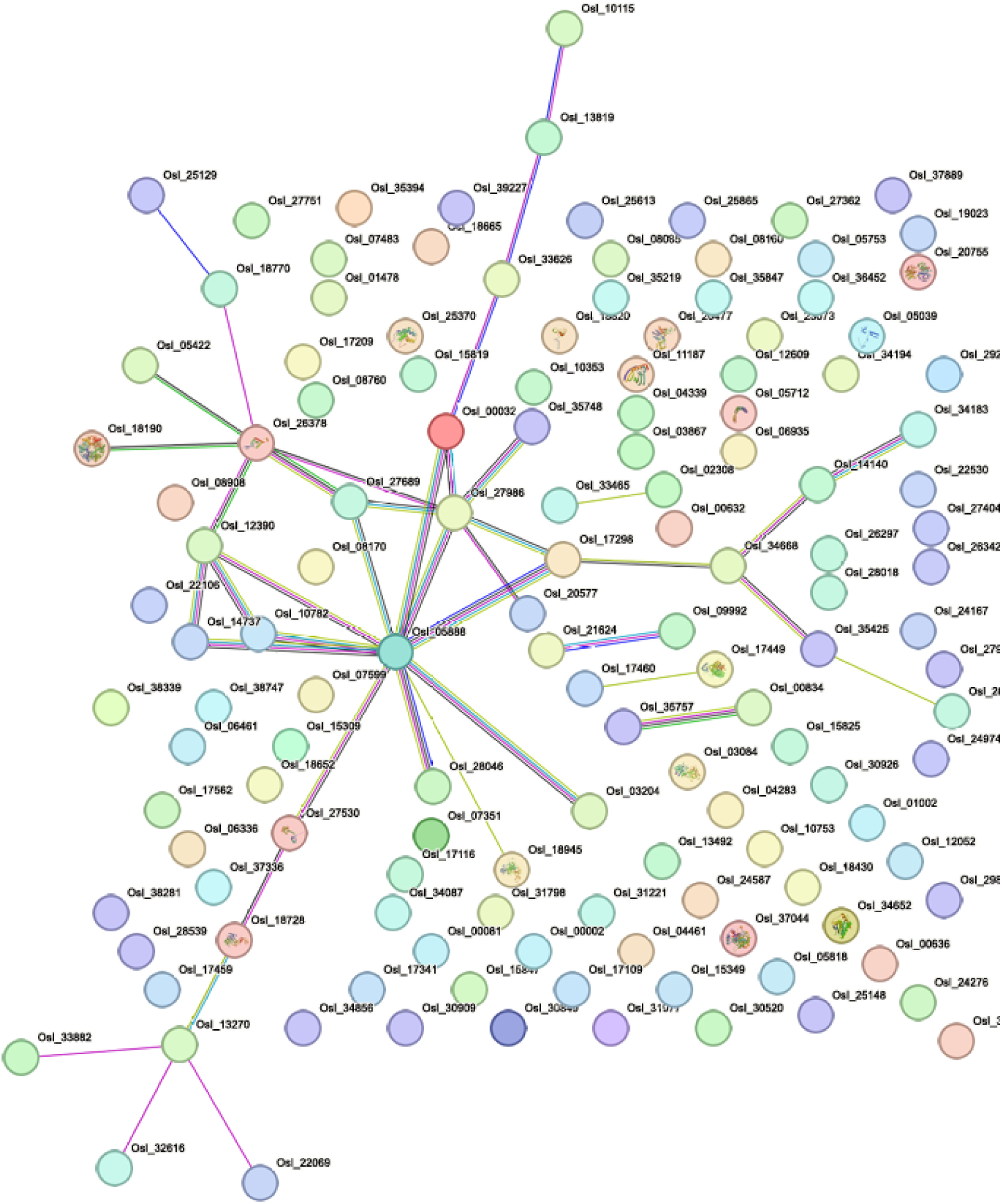
Association network of 134 identified PSNOs candidates enriching the *O. sativa-derived* S-nitrosoproteome. The nodes represent protein candidates, while edges represent PPIs. The empty nodes, in contrast with the filled nodes, represent proteins with an undetermined 3D structure.

## Notes

### Competing Interest Statement

The authors have declared no competing interest.

## Reference

1. Lindermayr C, Saalbach G, Durner J. Proteomic identification of S-nitrosylated proteins in Arabidopsis. Plant Physiol. 2005 Mar;137(3):921–30. doi: 10.1104/pp.104.058719.

2. Kolbert Z, Barroso JB, Brouquisse R, Corpas FJ, Gupta KJ, Lindermayr C, Loake GJ, Palma JM, Petřivalský M, Wendehenne D, Hancock JT. A forty year journey: The generation and roles of NO in plants. Nitric Oxide. 2019 Dec 1;93:53–70. doi: 10.1016/j.niox.2019.09.006.

3. Delledonne M, Zeier J, Marocco A, Lamb C. Signal interactions between nitric oxide and reactive oxygen intermediates in the plant hypersensitive disease resistance response. Proc Natl Acad Sci U S A. 2001 Nov 6;98(23):13454–9. doi: 10.1073/pnas.231178298.

4. Lindermayr C. Crosstalk between reactive oxygen species and nitric oxide in plants: Key role of S-nitrosoglutathione reductase. Free Radic Biol Med. 2018 Jul;122:110–115. doi: 10.1016/j.freeradbiomed.2017.11.027.

5. Zaffagnini M, De Mia M, Morisse S, Di Giacinto N, Marchand CH, Maes A, Lemaire SD, Trost P. Protein S-nitrosylation in photosynthetic organisms: A comprehensive overview with future perspectives. Biochim Biophys Acta. 2016 Aug;1864(8):952–66. doi: 10.1016/j.bbapap.2016.02.006.

6. Begara-Morales JC, Sánchez-Calvo B, Chaki M, Valderrama R, Mata-Pérez C, López-Jaramillo J, Padilla MN, Carreras A, Corpas FJ, Barroso JB. Dual regulation of cytosolic ascorbate peroxidase (APX) by tyrosine nitration and S-nitrosylation. J Exp Bot. 2014 Feb;65(2):527–38. doi: 10.1093/jxb/ert396.

7. Mata-Pérez C, Sánchez-Calvo B, Padilla MN, Begara-Morales JC, Luque F, Melguizo M, Jiménez-Ruiz J, Fierro-Risco J, Peñas-Sanjuán A, Valderrama R, Corpas FJ, Barroso JB. Nitro-Fatty Acids in Plant Signaling: Nitro-Linolenic Acid Induces the Molecular Chaperone Network in Arabidopsis. Plant Physiol. 2016 Feb;170(2):686–701. doi: 10.1104/pp.15.01671.

8. Arasimowicz-Jelonek M, Floryszak-Wieczorek J. A physiological perspective on targets of nitration in NO-based signaling networks in plants. J Exp Bot. 2019 Aug 29;70(17):4379–4389. doi: 10.1093/jxb/erz300.

9. Feechan A, Kwon E, Yun BW, Wang Y, Pallas JA, Loake GJ. A central role for S-nitrosothiols in plant disease resistance. Proc Natl Acad Sci U S A. 2005 May 31;102(22):8054–9. doi: 10.1073/pnas.0501456102.

10. Foresi N, Correa-Aragunde N, Parisi G, Caló G, Salerno G, Lamattina L. Characterization of a nitric oxide synthase from the plant kingdom: NO generation from the green alga Ostreococcus tauri is light irradiance and growth phase dependent. Plant Cell. 2010 Nov;22(11):3816–30. doi: 10.1105/tpc.109.073510.

11. Correa-Aragunde N, Nejamkin A, Del Castello F, Foresi N, Lamattina L. Nitric oxide synthases from photosynthetic organisms improve growth and confer nitrosative stress tolerance in E. coli. Insights on the pterin cofactor. Nitric Oxide. 2022 Feb 1;119:41–49. doi: 10.1016/j.niox.2021.12.005.

12. Tun NN, Santa-Catarina C, Begum T, Silveira V, Handro W, Floh EI, Scherer GF. Polyamines induce rapid biosynthesis of nitric oxide (NO) in Arabidopsis thaliana seedlings. Plant Cell Physiol. 2006 Mar;47(3):346–54. doi: 10.1093/pcp/pci252.

13. Desikan R, Griffiths R, Hancock J, Neill S. A new role for an old enzyme: nitrate reductase-mediated nitric oxide generation is required for abscisic acid-induced stomatal closure in Arabidopsis thaliana. Proc Natl Acad Sci U S A. 2002 Dec 10;99(25):16314–8. doi: 10.1073/pnas.252461999.

14. Wang BL, Tang XY, Cheng LY, Zhang AZ, Zhang WH, Zhang FS, Liu JQ, Cao Y, Allan DL, Vance CP, Shen JB. Nitric oxide is involved in phosphorus deficiency-induced cluster-root development and citrate exudation in white lupin. New Phytol. 2010 Sep;187(4):1112–1123. doi: 10.1111/j.1469-8137.2010.03323.x.

15. Romero-Puertas MC, Campostrini N, Mattè A, Righetti PG, Perazzolli M, Zolla L, Roepstorff P, Delledonne M. Proteomic analysis of S-nitrosylated proteins in Arabidopsis thaliana undergoing hypersensitive response. Proteomics. 2008 Apr;8(7):1459–69. doi: 10.1002/pmic.200700536.

16. Wawer I, Bucholc M, Astier J, Anielska-Mazur A, Dahan J, Kulik A, Wysłouch- Cieszynska A, Zareba-Kozioł M, Krzywinska E, Dadlez M, Dobrowolska G, Wendehenne D. Regulation of Nicotiana tabacum osmotic stress-activated protein kinase and its cellular partner GAPDH by nitric oxide in response to salinity. Biochem J. 2010 Jul 1;429(1):73–83. doi: 10.1042/BJ20100492.

17. Lamotte O, Courtois C, Dobrowolska G, Besson A, Pugin A, Wendehenne D. Mechanisms of nitric-oxide-induced increase of free cytosolic Ca2+ concentration in Nicotiana plumbaginifolia cells. Free Radic Biol Med. 2006 Apr 15;40(8):1369–76. doi: 10.1016/j.freeradbiomed.2005.12.006.

18. Camejo D, Romero-Puertas Mdel C, Rodríguez-Serrano M, Sandalio LM, Lázaro JJ, Jiménez A, Sevilla F. Salinity-induced changes in S-nitrosylation of pea mitochondrial proteins. J Proteomics. 2013 Feb 21;79:87–99. doi: 10.1016/j.jprot.2012.12.003.

19. de Pinto MC, Locato V, Sgobba A, Romero-Puertas Mdel C, Gadaleta C, Delledonne M, De Gara L. S-nitrosylation of ascorbate peroxidase is part of programmed cell death signaling in tobacco Bright Yellow-2 cells. Plant Physiol. 2013 Dec;163(4):1766–75. doi: 10.1104/pp.113.222703.

20. Jedelská T, Kraiczová VŠ, Berčíková L, Činčalová L, Luhová L, Petřivalský M. Tomato Root Growth Inhibition by Salinity and Cadmium Is Mediated By S-Nitrosative Modifications of ROS Metabolic Enzymes Controlled by S-Nitrosoglutathione Reductase. Biomolecules. 2019 Aug 21;9(9):393. doi: 10.3390/biom9090393.

21. Abat JK, Deswal R. Differential modulation of S-nitrosoproteome of Brassica juncea by low temperature: change in S-nitrosylation of Rubisco is responsible for the inactivation of its carboxylase activity. Proteomics. 2009 Sep;9(18):4368–80. doi: 10.1002/pmic.200800985.

22. Abat JK, Mattoo AK, Deswal R. S-nitrosylated proteins of a medicinal CAM plant Kalanchoe pinnata- ribulose-1,5-bisphosphate carboxylase/oxygenase activity targeted for inhibition. FEBS J. 2008 Jun;275(11):2862–72. doi: 10.1111/j.1742-4658.2008.06425.x.

23. Chaki M, Valderrama R, Fernández-Ocaña AM, Carreras A, Gómez-Rodríguez MV, Pedrajas JR, Begara-Morales JC, Sánchez-Calvo B, Luque F, Leterrier M, Corpas FJ, Barroso JB. Mechanical wounding induces a nitrosative stress by down-regulation of GSNO reductase and an increase in S-nitrosothiols in sunflower (Helianthus annuus) seedlings. J Exp Bot. 2011 Mar;62(6):1803–13. doi: 10.1093/jxb/erq358.

24. Cheng T, Chen J, Ef AA, Wang P, Wang G, Hu X, Shi J. Quantitative proteomics analysis reveals that S-nitrosoglutathione reductase (GSNOR) and nitric oxide signaling enhance poplar defense against chilling stress. Planta. 2015 Dec;242(6):1361–90. doi: 10.1007/s00425-015-2374-5.

25. Lin A, Wang Y, Tang J, Xue P, Li C, Liu L, Hu B, Yang F, Loake GJ, Chu C. Nitric oxide and protein S-nitrosylation are integral to hydrogen peroxide-induced leaf cell death in rice. Plant Physiol. 2012 Jan;158(1):451–64. doi: 10.1104/pp.111.184531.

26. Mun BG, Shahid M, Lee GS, Hussain A, Yun BW. A Novel RHS1 Locus in Rice Attributes Seed-Pod Shattering by the Regulation of Endogenous S-Nitrosothiols. Int J Mol Sci. 2022 Oct 30;23(21):13225. doi: 10.3390/ijms232113225.

27. Lehner C, Kerschbaum HH, Lütz-Meindl U. Nitric oxide suppresses growth and development in the unicellular green alga Micrasterias denticulata. J Plant Physiol. 2009 Jan 30;166(2):117–27. doi: 10.1016/j.jplph.2008.02.012.

28. Chatterji A, Sengupta R. Stability of S-nitrosothiols and S-nitrosylated proteins: A struggle for cellular existence! J Cell Biochem. 2021 Nov;122(11):1579–1593. doi: 10.1002/jcb.30139.

29. Das N, Bhattacharya S, Bhattacharyya S, Maiti MK. Identification of alternatively spliced transcripts of rice phytochelatin synthase 2 gene OsPCS2 involved in mitigation of cadmium and arsenic stresses. Plant Mol Biol. 2017 May;94(1-2):167–183. doi: 10.1007/s11103-017-0600-1.

30. Ren X, Sengupta R, Lu J, Lundberg JO, Holmgren A. Characterization of mammalian glutaredoxin isoforms as S-denitrosylases. FEBS Lett. 2019 Jul;593(14):1799–1806. doi: 10.1002/1873-3468.13454.

31. Mathews WR, Kerr SW. Biological activity of S-nitrosothiols: the role of nitric oxide. J Pharmacol Exp Ther. 1993 Dec;267(3):1529–37.

32. Eyer P, Podhradský D. Evaluation of the micromethod for determination of glutathione using enzymatic cycling and Ellman’s reagent. Anal Biochem. 1986 Feb 15;153(1):57–66. doi: 10.1016/0003-2697(86)90061-8.

33. Tietze F. Enzymic method for quantitative determination of nanogram amounts of total and oxidized glutathione: applications to mammalian blood and other tissues. Anal Biochem. 1969 Mar;27(3):502–22. doi: 10.1016/0003-2697(69)90064-5.

34. Samaddar S, Chakraborty S, Sengupta R, Ghosh S. Detection and proteomic identification of in vivo S-nitrosylated proteins in Vibrio cholerae: A novel evidence. Nitric Oxide. 2025 Sep 30;159:63–77. doi: 10.1016/j.niox.2025.09.005.

35. Stoyanovosky DA, Goldman R, Jonnalagadda SS, Day BW, Claycamp HG, Kagan VE. Detection and characterization of the electron paramagnetic resonance-silent glutathionyl- 5,5-dimethyl-1-pyrroline N-oxide adduct derived from redox cycling of phenoxyl radicals in model systems and HL-60 cells. Arch Biochem Biophys. 1996 Jun 1;330(1):3–11. doi: 10.1006/abbi.1996.0219.

36. Romero JM, Bizzozero OA. Intracellular glutathione mediates the denitrosylation of protein nitrosothiols in the rat spinal cord. J Neurosci Res. 2009 Feb 15;87(3):701–9. doi: 10.1002/jnr.21897.

37. Chakraborty S, Sircar E, Bhattacharyya C, Choudhuri A, Mishra A, Dutta S, Bhatta S, Sachin K, Sengupta R. S-Denitrosylation: A Crosstalk between Glutathione and Redoxin Systems. Antioxidants (Basel). 2022 Sep 28;11(10):1921. doi: 10.3390/antiox11101921.

38. Raj Rai S, Bhattacharyya C, Sarkar A, Chakraborty S, Sircar E, Dutta S, Sengupta R. Glutathione: Role in Oxidative/Nitrosative Stress, Antioxidant Defense, and Treatments. ChemistrySelect. 2021 May 14;6(18):4566–90. doi:10.1002/slct.202100773.

39. Zechmann B, Müller M. Subcellular compartmentation of glutathione in dicotyledonous plants. Protoplasma. 2010 Oct;246(1-4):15–24. doi: 10.1007/s00709-010-0111-2.

40. Zechmann B. Compartment-specific importance of glutathione during abiotic and biotic stress. Front Plant Sci. 2014 Oct 20;5:566. doi: 10.3389/fpls.2014.00566.

41. Koffler BE, Bloem E, Zellnig G, Zechmann B. High resolution imaging of subcellular glutathione concentrations by quantitative immunoelectron microscopy in different leaf areas of Arabidopsis. Micron. 2013 Feb;45:119–28. doi: 10.1016/j.micron.2012.11.006.

42. Gasperl A, Zellnig G, Kocsy G, Müller M. Organelle-specific localization of glutathione in plants grown under different light intensities and spectra. Histochem Cell Biol. 2022 Sep;158(3):213–227. doi: 10.1007/s00418-022-02103-2.

43. Airaki M, Sánchez-Moreno L, Leterrier M, Barroso JB, Palma JM, Corpas FJ. Detection and quantification of S-nitrosoglutathione (GSNO) in pepper (Capsicum annuum L.) plant organs by LC-ES/MS. Plant Cell Physiol. 2011 Nov;52(11):2006–15. doi: 10.1093/pcp/pcr133.

44. Mata-Pérez C, Padilla MN, Sánchez-Calvo B, Begara-Morales JC, Valderrama R, Chaki M, Aranda-Caño L, Moreno-González D, Molina-Díaz A, Barroso JB. Endogenous Biosynthesis of S-Nitrosoglutathione From Nitro-Fatty Acids in Plants. Front Plant Sci. 2020 Jun 30;11:962. doi: 10.3389/fpls.2020.00962.

45. Mata-Pérez C, Sánchez-Calvo B, Padilla MN, Begara-Morales JC, Valderrama R, Corpas FJ, Barroso JB. Nitro-fatty acids in plant signaling: New key mediators of nitric oxide metabolism. Redox Biol. 2017 Apr;11:554–561. doi: 10.1016/j.redox.2017.01.002.

46. Yamasaki H, Sakihama Y. Simultaneous production of nitric oxide and peroxynitrite by plant nitrate reductase: in vitro evidence for the NR-dependent formation of active nitrogen species. FEBS Lett. 2000 Feb 18;468(1):89–92. doi: 10.1016/s0014-5793(00)01203-5.

47. Rockel P, Strube F, Rockel A, Wildt J, Kaiser WM. Regulation of nitric oxide (NO) production by plant nitrate reductase in vivo and in vitro. J Exp Bot. 2002 Jan;53(366):103–10.

48. Sengupta R, Holmgren A. The role of thioredoxin in the regulation of cellular processes by S-nitrosylation. Biochim Biophys Acta. 2012 Jun;1820(6):689–700. doi: 10.1016/j.bbagen.2011.08.012.

49. Sakuma R, Nishina T, Kitamura M. Deproteinizing methods evaluated for determination of uric acid in serum by reversed-phase liquid chromatography with ultraviolet detection. Clin Chem. 1987 Aug;33(8):1427–30.

50. Stoyanovsky DA, Maeda A, Atkins JL, Kagan VE. Assessments of thiyl radicals in biosystems: difficulties and new applications. Anal Chem. 2011 Sep 1;83(17):6432–8. doi: 10.1021/ac200418s.

51. Sircar E, Stoyanovsky DA, Billiar TR, Holmgren A, Sengupta R. Analysis of glutathione mediated S-(de)nitrosylation in complex biological matrices by immuno-spin trapping and identification of two novel substrates. Nitric Oxide. 2022 Jan 1;118:26–30. doi: 10.1016/j.niox.2021.10.008.

